# Retinal morphology in *Astyanax mexicanus* during degeneration

**DOI:** 10.1101/706374

**Authors:** Amany Emam, Marina Yoffe, Henry Cardona, Daphne Soares

## Abstract

The teleost *Astyanax mexicanus* is extant in two readily available forms. One that lives in Mexican rivers and various convergent forms that live in nearby caves. These fish are born with eyes but in the cavefish they degenerate during development. It is known that the lens of cavefish undergoes apoptosis and that some cells in the neuroretina also die. It has not been described, however, if glia and various components of the neuroretina form before complete eye degeneration. Here we examined the development of the retina of the closest living ancestor that lives in the rivers and members of two lineages of cavefish. We report that although the neuroretina is smaller and more compact, it has all cell types and layers including amacrine cells and Muller glia. While various makers for photoreceptors are present in the cavefish inner segments, the outer segments of the photoreceptors in cavefish are missing from the earliest stages examined. This shows that the machinery for visual transducing discs might still be present but not organized in one part of the cell. It is interesting to note that the deficiencies in Astyanax cavefish resemble retinal diseases, such as retinitis pigmentosa.

## Introduction

### Cavefishes

One of the hallmarks of troglobitic species is the absence of external visual structures. Darkness has led to the degeneration of the visual system and as animals become more cave adapted their eyes become smaller and eventually vanish completely. This gradual loss is true in all troglobites, may they be invertebrates, salamanders or fishes (Barr, 1968; Fong et al., 1995; Protas and Jeffery, 2012; Culver and Pipan, 2016). The Mexican tetra, *Astyanax mexicanus*, provides a particularly good window into how adaptive changes of visual networks occured in response to a life in darkness. This teleost is undergoing allopatric speciation and is extant in two readily available forms: an ancestral, river-dwelling form (surface fish) and various derived, cave-dwelling forms (each one named after the cave in which they are endemic). Mitochondrial DNA sequences, as well as morphological differences, suggest that cave forms are divided into two lineages. Both lineages contain eyeless fish populations, lending support to the idea that eye loss has evolved various times. One lineage (A) includes *Astyanax* cavefishes from the Subterráneo, Pachón and Chica populations, as well as surrounding surface fish (Dowling et al., 2002; Porter at al., 2007; Gross, 2012; Bradic et al., 2013). A second lineage (B) contains only cavefish from Los Sabinos, Curva and Tinaja, which are morphologically distinct in that they are anteroposteriorially compressed having one less set of ribs, and are not related to nearby epigean populations (Dowling et al., 2002). Studies in *Astyanax* have shown that although adult cavefish lack eyes, eye development is nevertheless initiated during embryogenesis (reviewed in Jeffery, 2001 and 2009). The eye is formed in the early embryo but later arrests, and degenerates, consequently sinking into the orbit. The lens itself undergoes apoptosis (Yamamoto and Jeffery, 2000) but since the eye is a developmental organizer for the forebrain its emergence remains under evolutionary constraint until the forebrain is at least partially formed, then it is released from adaptive pressure and degenerates (Retaux et al., 2008).

### Retinogenesis

The vertebrate retina itself is remarkably conserved for over 500 MY (Lamb et al., 2007). Retinal histogenesis is basically a process of delamination of the neuroepithelium. At early stages, the primary optic primordia create outpockets from the forebrain that arrive at the surface ectoderm, then invaginate to create an optic cup. The delamination comes when the inner surface of the cup becomes the sensory portion of the retina while the outer surfaces develops into the retinal pigmented epithelium (Chow and Lang, 2001). The retina contains six neuronal types and one glia. These are distributed in organized fashion in three nuclear layers and two plexiform layers (Livesey and Cepko, 2001), parallel to the surface of the eye. Cone and rod photoreceptors are located in the external most nuclear layer, where they act and sensory transducers. Retinal ganglion cells (RGCs) are located in the innermost region, closest to the lens. These are the only cells that send projections from the eye to innervate the visual part of the fish brain, the optic tectum (OT). The other neurons in the retina, bipolar, horizontal and amacrine cells form the inner nuclear layer (INL), located between the outer nuclear layer (ONL, cell bodies of photoreceptors) and the ganglion cell layer (GCL, cell bodies of RGCs). While bipolar cells connect photoreceptors to the RGCs in the vertical axis, amacrine cells transform the visual signal by integrating information in a horizontal plane. There are reports however, of displaced amacrine cells in the GCL in mammals (Perry and Walker, 1980, Wong and Hughes, 1987). Horizontal cells are connected to photoreceptors and mediate lateral inhibition while amacrine cells connect RGCs to modulate the visual signal further. Muller cells are elongated glial cells that vertically span the entirety of neuroretina layers (Newman and Reichenbach, 1996), and support the retina structurately and functionately. All seven cell types arise from the delamination of the cell marginal zone (CMZ) located on the rim on the retina, near the lens. In the mammalian retina RGCs and horizontal cells delaminate first, followed by cones, amacrine, rods, bipolar and finally Muller glial cells (Sidman, 1960; LaVail et al., 1991; Carter-Dawson and LaVail, 1979). In teleosts there is a delay between the emergence of rods and cones and in some species the larval retina has cones but no rods (Blaxter and Staines, 1970; Blaxter, 1975). On the other hand, in many species of fish the production of rods continues into adulthood (Muller, 1952; Munk and Jørgensen, 1983; Raymond, 1985; Fernald, 1989; Evans and Fernald, 1993).

Here we examine the development of the retina of *Astyanax* from both lineages using immunohistochemistry during the period in which the eye is developing but the lens is degenerating. We also compared them to the closest living surface fish ancestor. We show that the basic organization of the neuroretina resembles the ancestral form, including all neuron types and Muller cells. However, the outer segments of the photoreceptors is absent and the outer nuclear layer is disorganized in both lineages of cavefish.

## Methods

Fish larvae originated from a breeding colony kept in a 12:12 light and dark cycle. Breeding was induced by gradually increasing the water temperature from 20°C to 25°C over two days, every two weeks. Fish larvae were used at 3 days post fertilization (dpf) and at 7 dpf, when the mouth appears and the fish start feeding.

For overall anatomy live larvae were imaged using a Nikon SMZ25 microscope equipped with NIS-elements D software. Standard length of the lavae was measured from head to tail, not including fin, results are given in mm with standard deviation. Animals were fixed in 4% Paraformaldehyde in 0.1M PB and maintained at 4°C at least overnight. Larva we cryoprotected in 30% Sucrose in 0.1M PBS before sectioning. Larva were embedded in Tissue-Tek^®^ O.C.T. compound medium and frozen at −80°C for a few minutes before transferred to a −20° Cryostat (ThermoFisher HM525NX). For cresyl violet sections, fish heads were cut at 10 micrometers and placed on chrome alum subbed slides. Slides were left overnight to dry and stained with cresyl violet for 20 minutes then briefly washed in reverse osmosis water. Slides were then dehydrated in an alcohol series and transferred into Histoclear. Finally, slides were mounted with permount media (Sigma) and left to dry overnight. Slides were imaged using Olympus PX50 microscope and Imaging software (Toup View).

For immunohistochemistry, 20 micrometer thick sections were cut and mounted onto chrome alum subbed glass slides. Slides were left to dry overnight at room temperature to ensure adhesion. Slides were washed in 1x PBS then blocked with normal goat serum (0.2% fetal bovine serum, 0.3% TritonX 100, 10% normal goat serum in PBS). Slides were incubated with primary antibodies overnight in a humid chamber using the following dilutions: Glial Fibrillary Acidic Protein (GFAP 1:100, Rabbit polyclonal, GeneTex GTX128741); Tyrosine hydroxylase (TH 1:100, mouse monoclonal, Millipore MAB318); Zinc finger protein 1 (ZPR-1 1:100, mouse monoclonal, Abcam ab174435); ZPR-3 (1:100 mouse monoclonal, ZIRC zpr3); Rhodopsin (1D4 1:100, mouse monoclonal Abcam ab5417); Calretinin (1:200, rabbit polyclonal, Abcam ab702); synaptic vesicle 2 (SV2 1:250 mouse monoclonal, DSHB AB2315387). After room temperature washes, slides were incubated in fluorescent secondary antibodies (1:500 Alexa Fluor 568, abcam, ab175471). To counterstain we sometimes used phalloidin 488 (1:25, Thermofisher A12379) and DAPI (1:5000, ThermoFisher Scientific, D3571). Slides were mounted in Fluoromount Aqueous Mounting Medium (Sigma F4680) and sealed with nail polish. Fluorescent images were captured in a SP8 confocal (Leica, USA) and processed using LAS X software (Leica).

For transmission electron microscopy, 18 larva of the two different ages were fixed whole in Karnovsky’s fixative (2.5% glutaraldehyde and 2% paraformaldehyde solution diluted in 0.1 M cacodylate buffer at pH 7.4; EMS RT15720; Karnovsky, 1965) for at least 24 hours at 15-20 °C and at a pH of 7.2. Larva were transferred to core imaging lab at the Robert Wood Johnson Medical School were specimens were washed in either cacodylate or phosphate buffer, and then treated with a 1 % osmium tetroxide fixative for 1 hour at room temperature. They were subsequently washed in maleate buffer (pH 6.0) and stained on block with a 1 % uranyl acetate solution in maleate buffer of pH 5.2. After serial dehydration in ethanol series, passed through propyleneoxide, and embedded in Epon. Ultrathin sections, cut serially with diamond knives on a Reichert-Jung Ultracut-E ultramicrotome, and stained with aqueous lead citrate. Sections were mounted either on a 200-mesh grid or on a 0.4 × 0.2 mm slot grid, gridswere examined with a JEOL 1200EX electron microscope with AMT-XR41 digital camera operated at 80 Kv. Images were compiled using LAS X, Photoshop CC 2019 and Illustrator CC 2019. We used Fiji (Schindelin et al., 2012) to measure the size of the fish and larvae and compiled the data using Excel (Microsoft). Images were pseudo colored for ease of interpretation to yellow instead of blue DAPI 461 nm, Magenta instead of green 488 nm and cyan instead of red 594 nm. Staining of ocular muscles with phalloidin were over exposed on purpose.

## Results

We aimed to visualize various aspects of the retina to determine what aspects of the circuitry were present during degeneration and which degenerated first. We mesured the size of the eye and used Cresyl violet to determine its gross morphology. We then used multiple antibodies to visualize the layers and components of the retina. Some features were confirmed at higher resolution provided by transmission electron microscopy.

### Fish and eye size

Fish lengths and eye sizes were measured at both developmental stages (Figure 1; n=10 per species for all measurements). At 3 dpf, surface fish (SF) length was 37.1±1.6 mm, Tinaja was 37.4 (±1.4) and Pachon was 37.4 (±1.4). At 7 dpf SF length was 42.0 (±1.9), Tinaja was 44.3 (±1.8) and was Pachon 45.4 (±1.3). Eye sizes at 3dpf for SF were 2.9 (±0.3), Tinaja were 1.9 (±0.2) and Pachon were 1.8 (± 1.1); at 7 dpf SF were 3.3 (±0.3), Tinaja were 1.3 (±0.3) and Pachon were 1.5 (±0.1). The ratio of body length to eye diameter at 3dpf was: SF 12.8, Tinaja 20.0 and Pachon 21.3; and at 7dpf to: SF 12.7, Tinaja 33.0 and Pachon 29.8.

**Figure 1.**
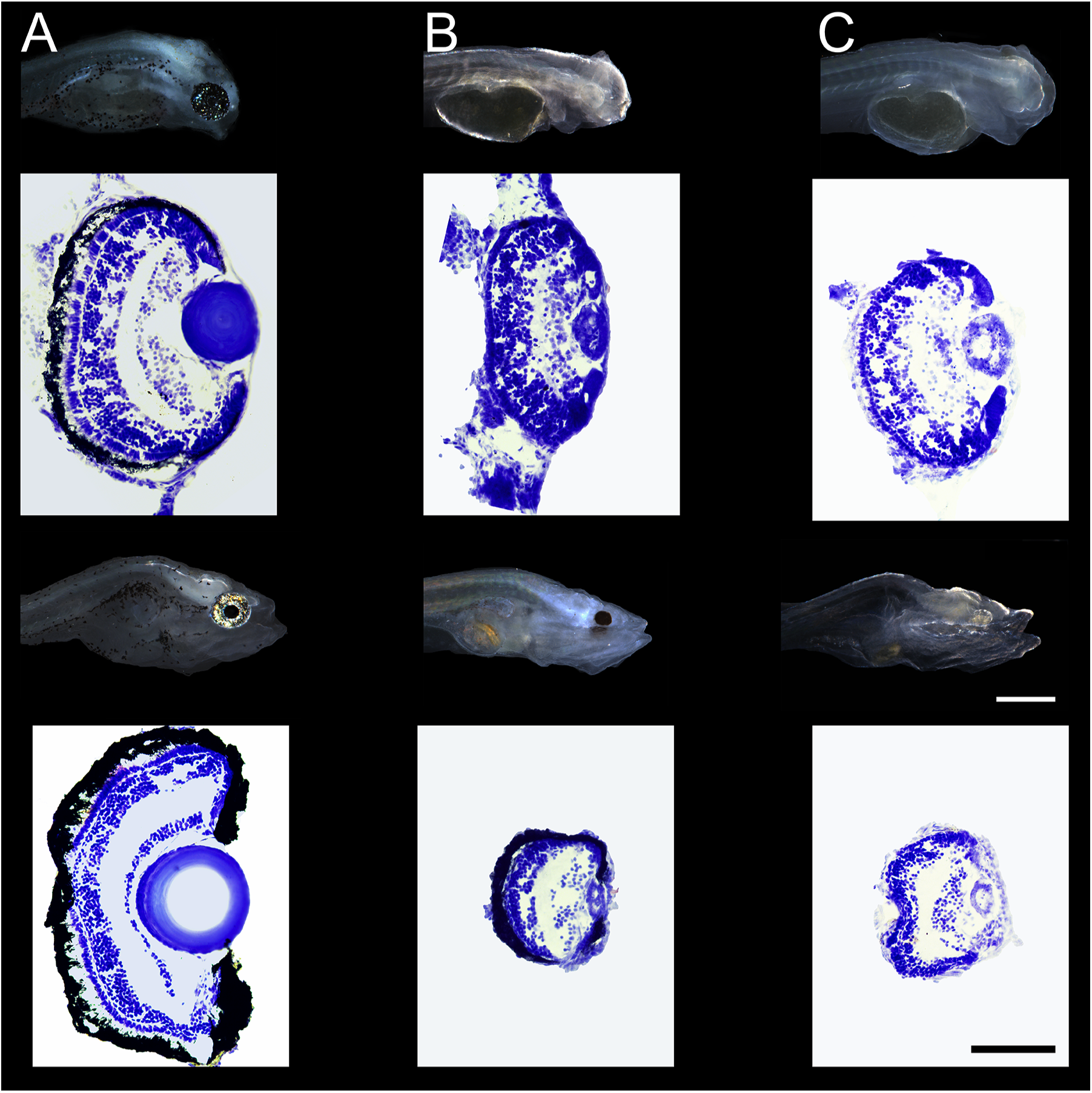
Larvae anatomy and gross histology. Whole mount and cresyl violet series. Top row 3 dpf and bottom row 7 dpf larva. A) Surface fish, B) Tinaja and C) Pachon. Scale bars are .05 mm for larvae images and 100 um for cresyl violet sections.

### Overall eye histology

In cresyl violet stained serial sections of eyes, 3 dpf SF had five well-organized retinal layers, with a prominent ciliary marginal zone (CMZ) and a pigmented layer (Figure 1). The retinas were less laminated in the cavefish. Tinaja showed a thick INL and sparse RGC layers but a thin IPL. The same was true for Pachon but the RGC was more diffuse. At 7dpf the SF showed an adult-like mature developmental pattern with a layered retina and a large lens. SF eye increased in overall size while cavefish eyes decreased in overall size and became less organized. The IPL of Tinaja had thickened and the RGC was thick with diffuse cell bodies. The ONL was compact but it was difficult to locate the OPL or the photoreceptor layers, although some cells were transverse to the plane of radiation. These cells could belong to the retinal pigmented epithelium (RPE), but were lateral to the pigmented layer. Pachon did not have a pigmented layer but followed the same pattern as Tinaja. The lens in the SF increased in size with age, while in both cavefish the lens appeared smaller as the fish grew older.

### Muller cells

Glial Fibrillary Acidic Protein (GFAP) is a marker for type III intermediate filament in Muller (glial) cells and was seen in both 3dpf and 7 dpf of all larvae (Figure 2). The SF had distinctive thinning projections from the RGC layer all the way to the photoreceptors, extending radially. Towards the lens the Muller cells had triangular shapes. In Tinja and Pachon at 3dpf GFAP processes were disorganized and did not pass the IPL. In Tinaja the processes were vertical to the retina and parallel to each other with processes intermingled at the entire length. Pachon 3dpf was the most disorganized, with processes extending to all directions although the bases were clearly seen towards the center of the eye. At 7dpf Muller cells were oriented radially in all species, including in Pachon, Triangular pedicles and thin processes extended into the photoreceptor layer in all fish.

**Figure 2.**
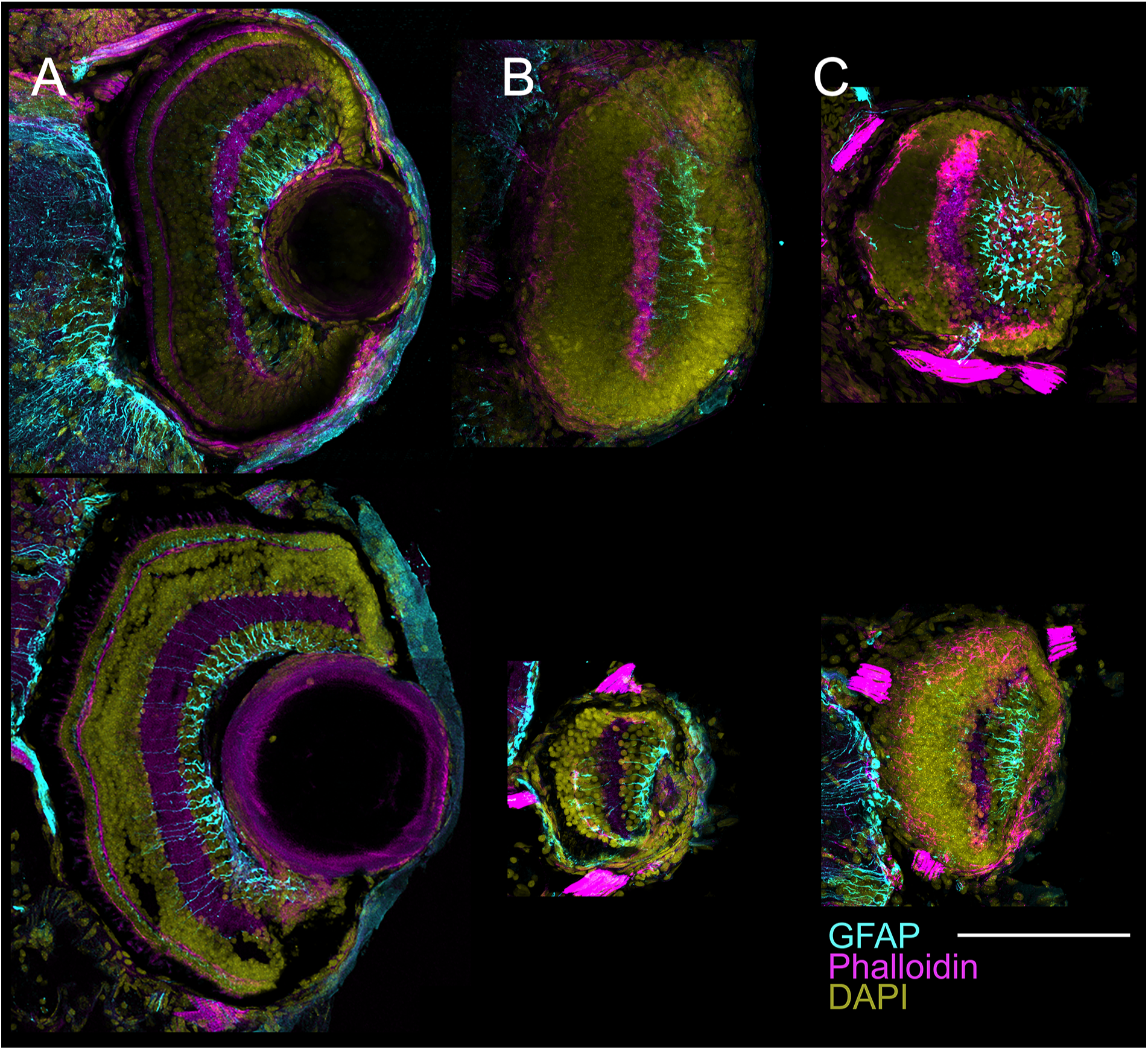
Muller cells are present in all fishes but disorganized in cavefish. Immunostain for Glial Fibrillary Acidic Protein (GFAP) reveals location of Muller cells in A) Surface fish, B) Tinaja and C) Pachon (top row 3 dpf and bottom row 7 dpf larva). Notice that in 3dpf in both cavefishes the cells are disorganized compared to the 7dpf. Scale bar 100 um.

### Dopaminergic neurons

Tyrosine Hydroxylase (TH) is the rate-limiting enzyme for dopamine biosynthesis and very few TH immunoreactive (TH-ir) cells were present in all fish at all stages (Figure 3) in the inner nuclear layer (INL). Somata were round to oval with projections extending to the inner plexiform layer (IPL). In the SF TH-ir puncta were also seen in the IPL and the outer plexiform layer (OPL), where the pedicles of the photoreceptors are located, at both developmental stages. At 3 and 7 dpf, immunoreactive cells were present in Tinaja and Pachon, yet puncta were only present in the IPL. At 7dpf some TH-ir cell bodies present in the IPL, only in the cavefishes. The cell exhibited a partial stellate appearance and gave rise to two or more primary dendrites with processes that were lightly branched and varicose, with terminal branches sometimes appearing very thin and punctate. By their morphology and immunoreactivity these cells could be amacrine and displaced amacrine cells.

**Figure 3.**
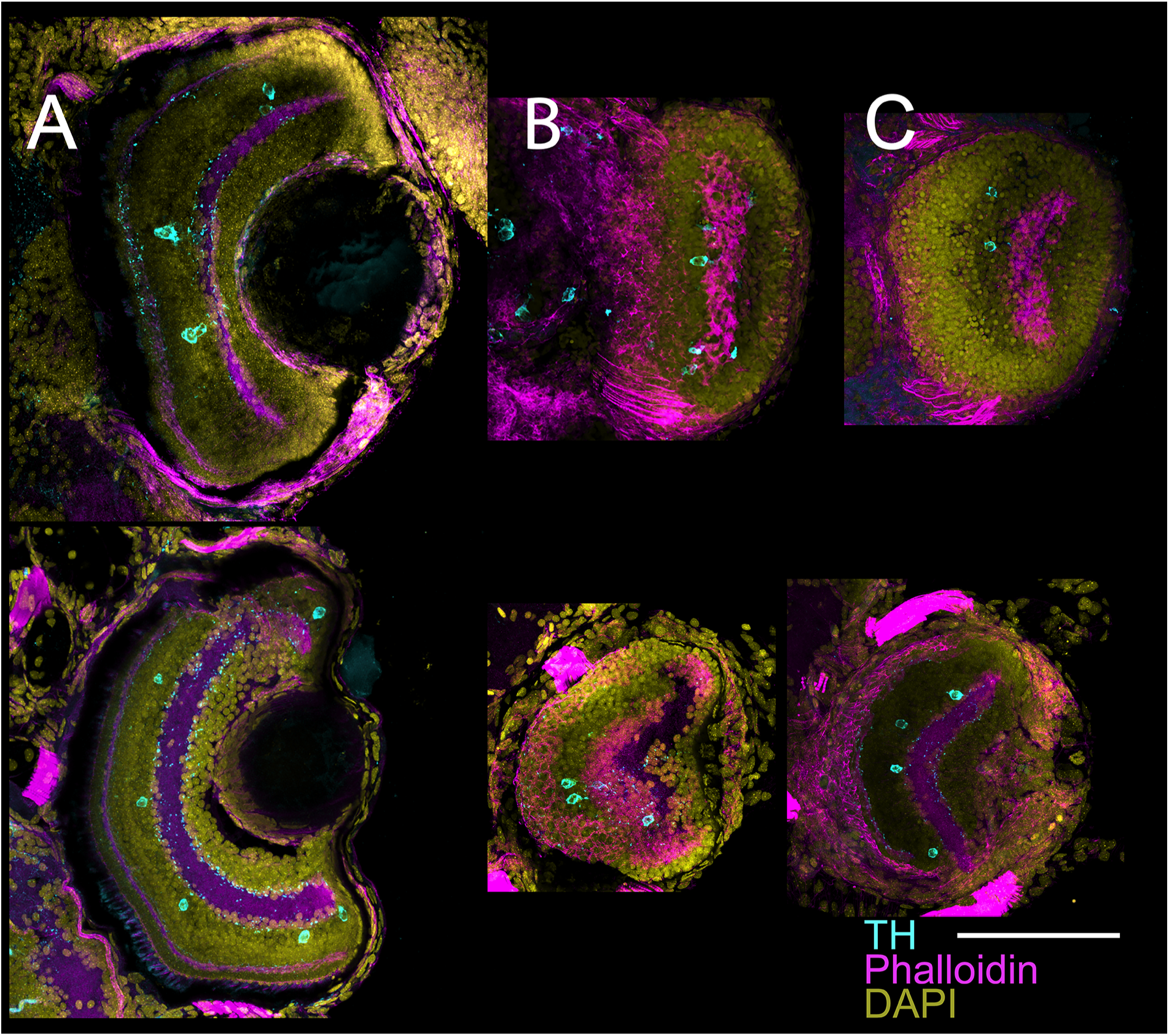
Dopaminergic positive amacrine cells and dispaced amacrine cells are sparse in the INL and GCL. Tyrosine Hydroxylase. Top row shows 3 dpf and bottom row show 7 dpf larva. A) Surface fish, B) Tinaja and C) Pachon. Scattered cell bodies are seen in the INL in cyan. Scale bar 100 um.

### Developing retinal ganglion cells and INL amacrine cells

Calretinin antibodies are markers for a calcium buffering protein and in the eye they stains a type of Amacrine cells with cell bodies in the inner nuclear layer (INL) and the GCL during development (Figure 4). Cells were clearly seen in all 3 and 7 dpf fish, although 3 dpf fish had less Calretinin-ir cells. In SF there were puncta in the IPL. The outer segments of photoreceptors were also lightly stained and in 7 dpf fish different retinal layers are more clearly defined in the SF. Tinaja at 7dpf showed no staining in the outer portions of the retina, where photoreceptors are located, while Pachon had calretinin positive diffuse staining in this area. This region could be the Retinal Pigment Epithelial since the cells bodies were oriented at parallel to the neural retina plane of organization.

**Figure 4.**
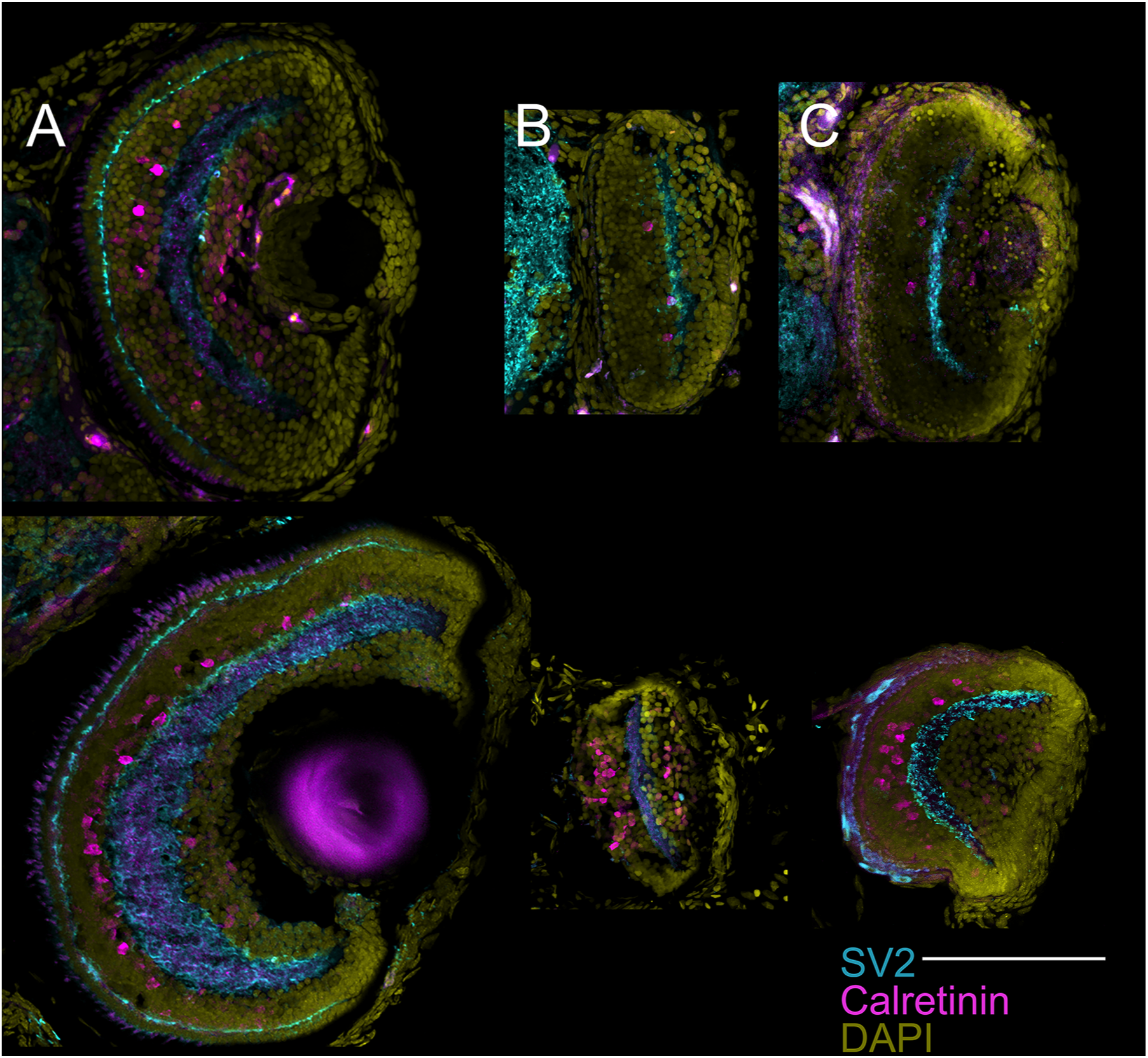
Developing retinal ganglion cells and INL amacrine cells. Immunostain for SV2 (cyan) and Calretenin (magenta) in A) Surface fish, B) Tinaja and C) Pachon at 3 (top) and 7 dpf (bottom) larva.. SV2 staining is most prevalent in the plexiform layers (OPL and IPL) while calretinin labels scattered amacrine cells. Scale bar 100 um.

### Chemical Synapses

Synaptic vesicle glycoprotein 2a (SV2) is a molecular component of chemical synapses that are expected in the plexiform layers. Indeed, SV2 immunoreactivity was located in the OPL and IPL (Figure 4) in the SF at both stages. In contrast, 3 dpf cavefish had no staining in the OPL but strong immunoreactivity in the IPL. At 7dpf many more puncta were seen in all fish. Only in 7 dpf Pachon fish, cells in the presumptive RPE also stained for SV2, but the staining pattern was not punctate as expected for synapses. Instead, diffuse staining outlined cell bodies (Figure 4 C, bottom).

### Cytoskeleton

Phalloidin stains for actin in the cytoskeleton of cells. Staining occurred in the outer segments of photoreceptors in the SF as well at in the OPL where the synaptic terminals of photoreceptors, as well as the presynaptic regions of the horizontal cells are located (Figure 5). Staining was similar between 3dpf and 7dpf so only 7 dpf is shown. There was a ring of acting below the outer segment layer. Presumably outer segment staining was labeling the photoreceptor connecting cilium, which connects the inner and outer segments. In both cavefish there was no staining in a clear outer segment layer but there were strong staining in a disorganized manner in what appears to be the OPL of Pachon. Actin filaments are noticeable surrounding a dark circular region which appear to be the somata of cells.

**Figure 5.**
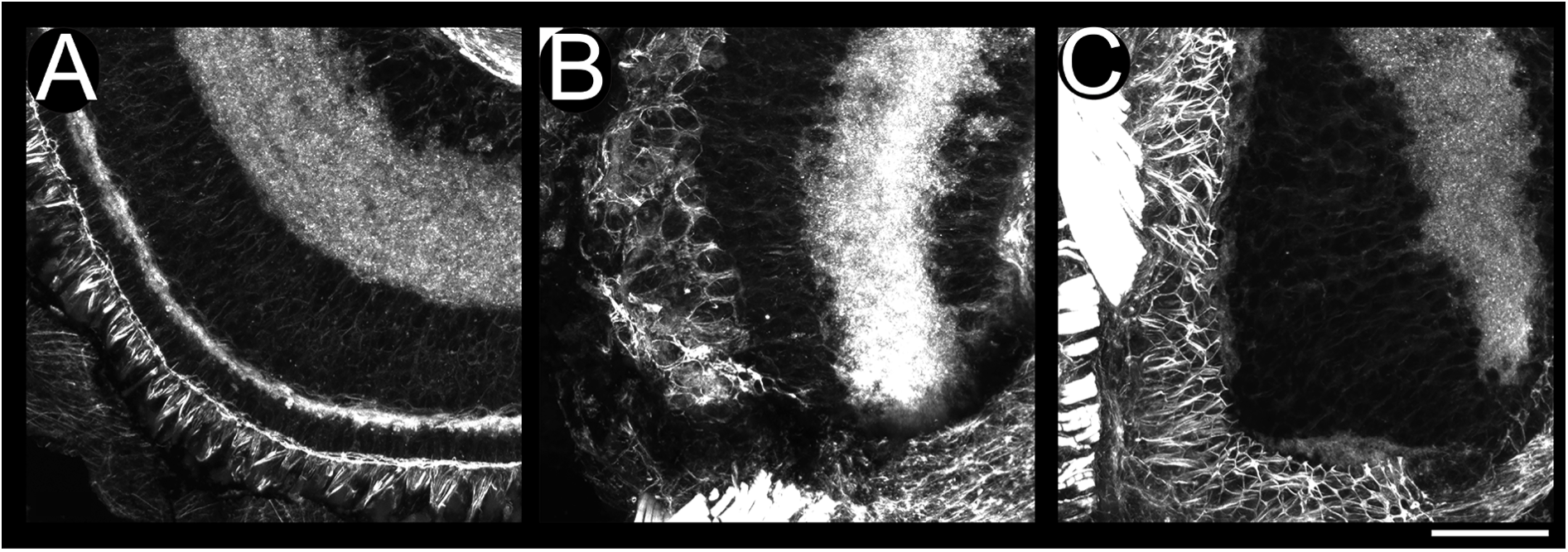
Actin filaments create a structural framework for the retina that is disorganized in the cavefish. 7 dpf staining with Phalloidin. A) Surface fish showing organized layers. B) Tinaja and C) Pachon showing disorganized layers with cytoskeleton in the cell bodies. Note what it seems to be the clia in Pachon and the optic nerve (overexposed). Scale bar 25 um.

### Long double cone photoreceptors

Rhodopsin 1D4 labels outer segments of long double cones but not rod photoreceptors (Figure 6). At 3 dpf signal could only be seen in the SF and neither cavefish had rhodopsin-ir. This signal only got stronger in the 7dpf SF, and the immunoreactivity could be seen in a patchy manner on the outer layers of the photoreceptors. The the complementary phalloidin staining showed actin prevalent in ocular muscle and plexiform layers. In the OPL, which was intertwined with Rhodopsin. The cavefishes also both showed Rhodopsin-ir intermingled with phalloidin staining

**Figure 6.**
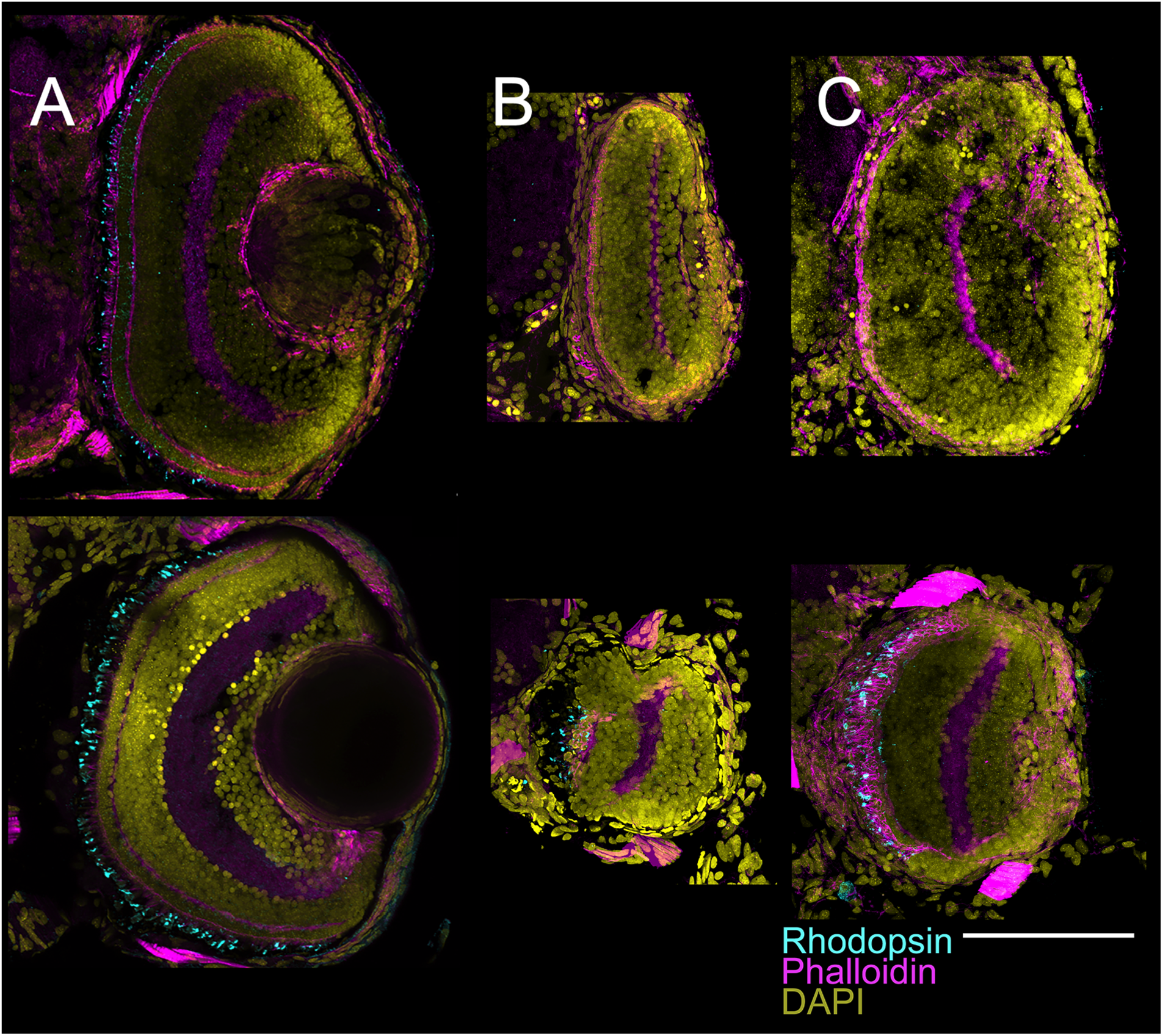
Photoreceptors show Rhodopsin in the inner segments of cavefish at 7pf. Rhodopsin. Top row shows 3 dpf and bottom row show 7 dpf larva. A) Surface fish, B) Tinaja and C) Pachon. At 3 dpf rhodopsin (cyan) is present in the outer segments in the SF, but not present in both cavefish forms. At 7 dpf signal appears in the cavefish. Note phalloidin staining of the photoreceptor layer. Scale bar 100 um.

### Green and red double cones

Zinc Finger Protein 1 stains arrestin 3a green and red double cones. We did not use phalloidin these sections for clarity. In 3dpf SF staining was seen mostly in the outer segments of the photoreceptors and in the pedicule areas. Scattered staining was seen in round cells in both cavefishes in a patchy manner, and the presumptive inner segment was clearly seen. At 7dpf, the SF immunoreactivity included the inner segments of the photoreceptors. Cells in both cavefishes were present with immunoreactivity in rounded cells. Clear nuclei were also seen. Unlike the SF, the cells were not stacked on an organized way and instead they were tangled on top of each other. (Figures 7, 9A and B).

**Figure 7.**
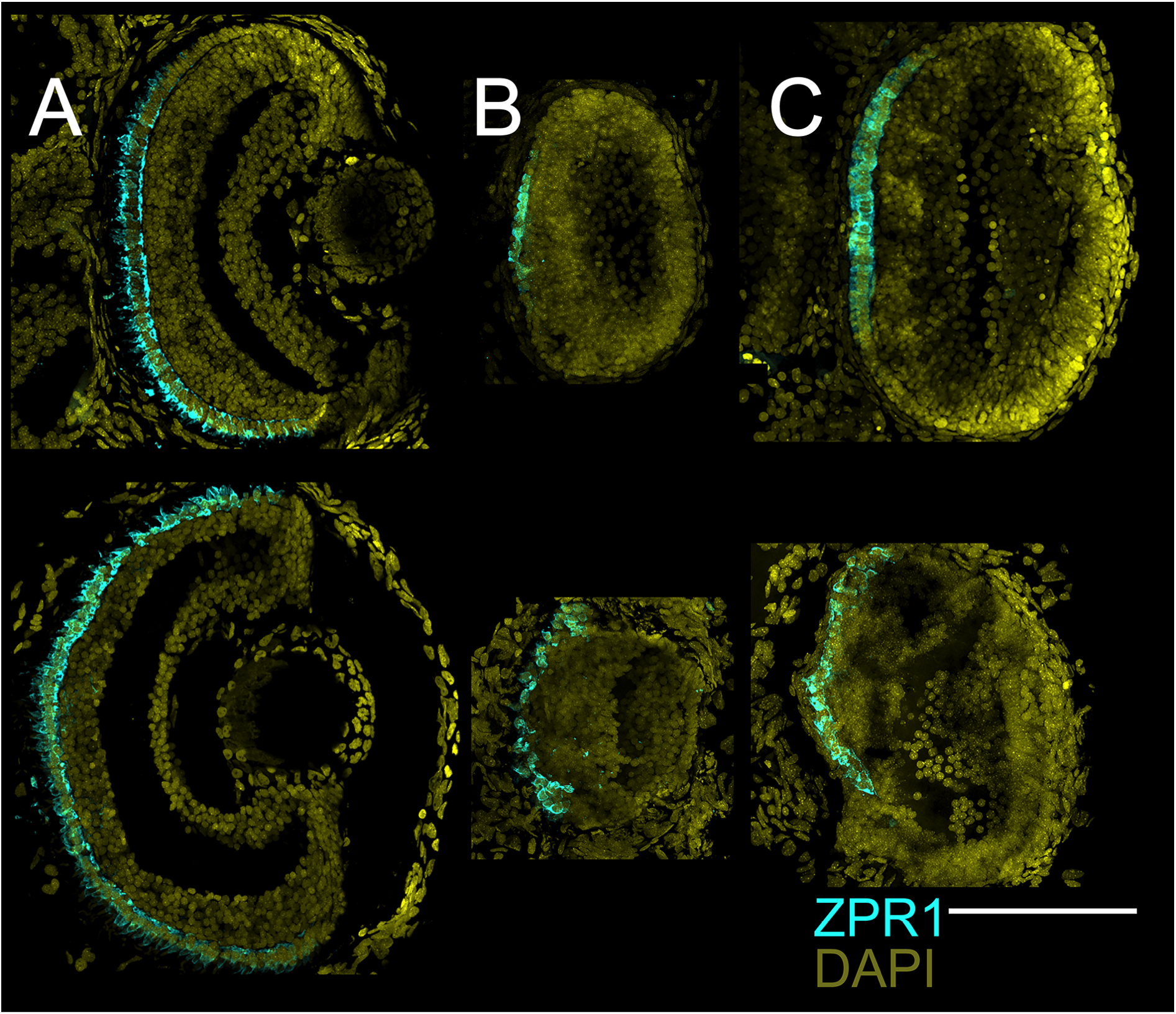
Inner segments of red and green double cones are present as early as 3dpf. ZPR1ir in photoreceptors. Top row shows 3 dpf and bottom row show 7 dpf larva. A) Surface fish, B) Tinaja and C) Pachon. In 3dpf ZPR1 is seen in photoreceptors in the SF and has patchy staining in the cavefish. At 7 dpf this pattern continues. Scale bar 100 um.

### Rods

Zinc Finger Protein 3 stains the outer segments of rod photoreceptors. In the 3dpf SF a clear layer of staining could be seen on the outer part of the photoreceptors (Figure 7). There was no signal in 3dpf Tinaja and in Pachon there were random small pockets of signal throughout the eye. At 7 dpf ZPR3 appears more layered in the SF with clear lamination. At this stage there was signal in Tinaja and Pachon, although it was dispersed throughout the outermost layers of the eye. Small portions of cells and puncta with clear outlines of the nuclei were stained. In both fish there were puncta like staining all over the outer layers of the eye. (Figures 8, 9C and D)

**Figure 8.**
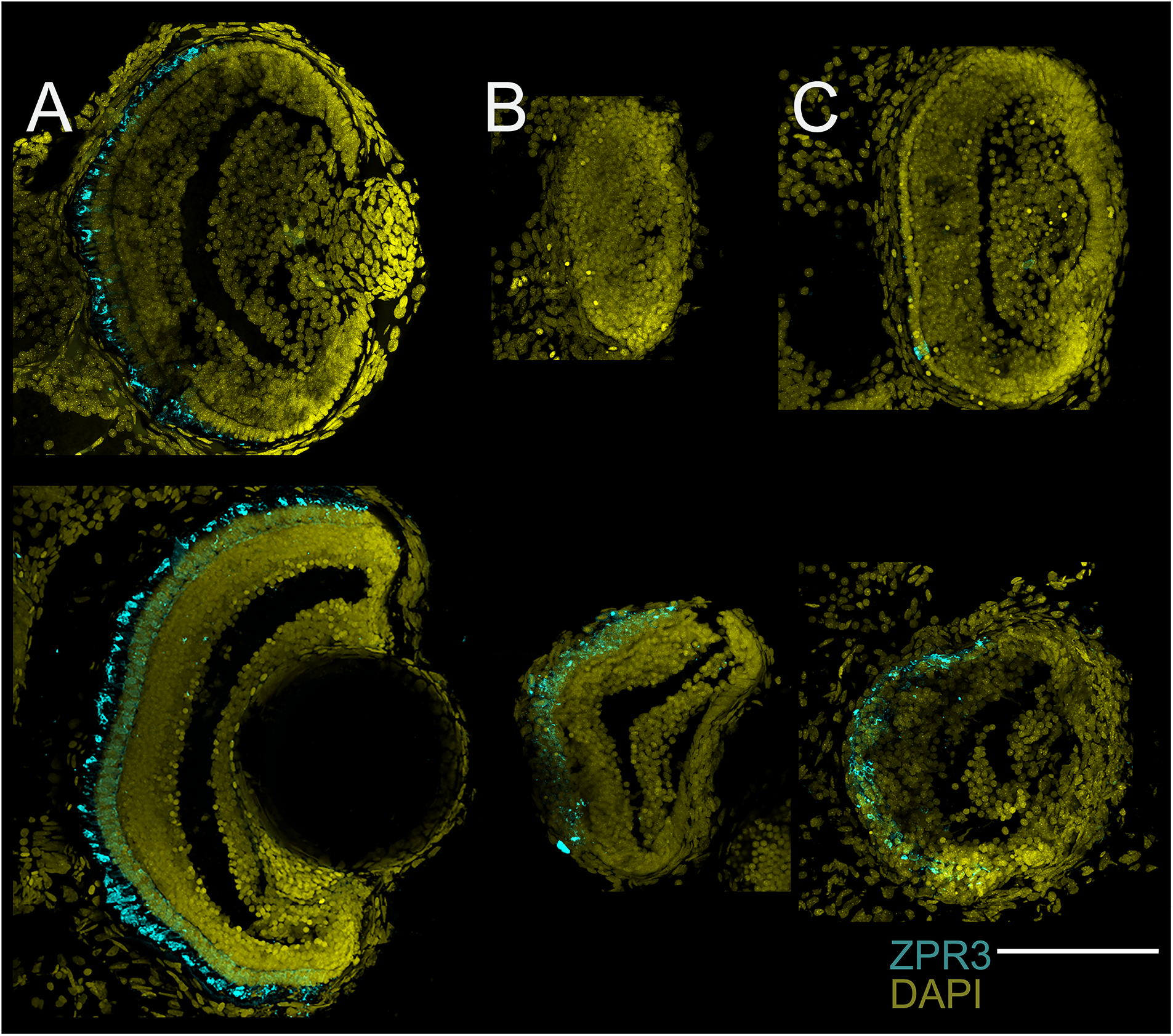
Rod photoreceptors appear later than cones in 7 dpf fish. Top row shows 3 dpf and bottom row show 7 dpf larva. A) Surface fish, B) Tinaja and C) Pachon. At 3 dpf ZPR3 is not seen in Tinaja, has strong ir in SF and very sparse staining in Pachon in what appears to be the inner segments. At 7dpf labeling is present in all fish forms. Scale 100 um

**Figure 9.**
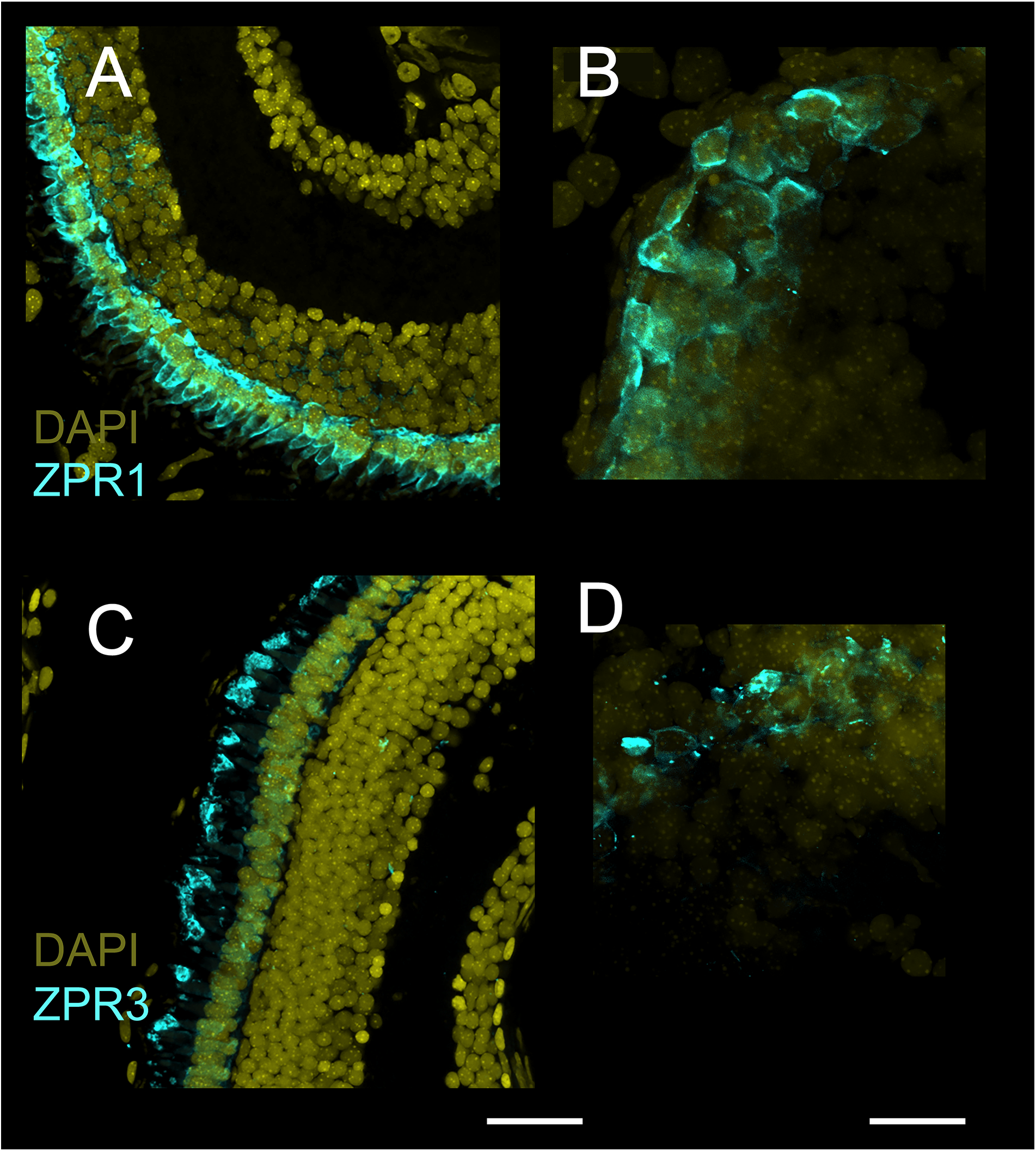
Cones and Rods are present in the 7dpf fish. A) surface fish ZPR1, B) Pachon ZPR1, C) surface fish ZPR3 and D) Pachon ZPR3. Scale bars 25 um for surface fish and 15 um for Pachon.

### Fine morphology of photoreceptors

Transmission Electron Microscopy provide subcellular resolution that allowed us to explore the structure of photoreceptor and in particularly their inner and outer segments. The most striking observation is that both cavefishes were missing the outer segment while the inner segments of the photoreceptors were present. In all fishes, the level of delamination in the photoreceptor layer increased with age. At 3dpf the SF already showed a layered retina with mature-like photoreceptors (not shown). Rods and Cones could be clearly seen. At 7 dpf this pattern became more refined and the retina looked adult-like (Figure 10). Both cavefishes, on the other hand, had distinct OPL, IPL and INL but the outer layers were difficult to identify, with Pachon more disorganized than Tinaja. Pigmentation in the RPE could be seen in the SF and Tinaja but not Pachon. The structures were smaller in Tinaja. structure and what we are looking at] Cells in INL that appeared to be apoptotic could be seen in both cavefish.

**Figure 10.**
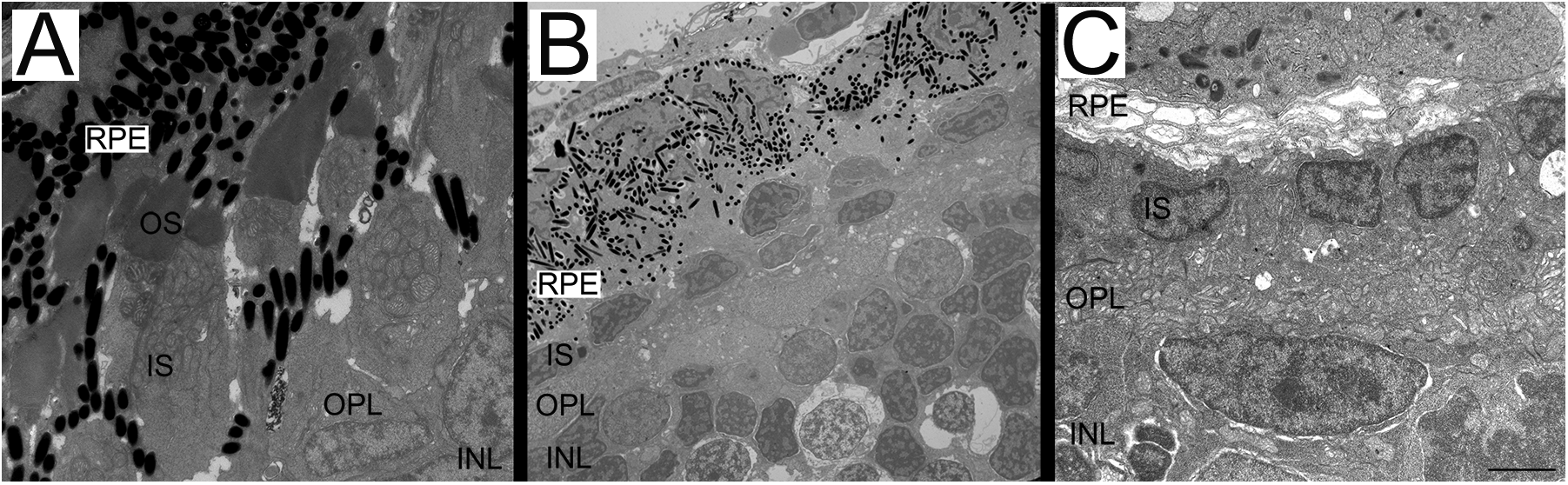
Transmission electron micrographs of 7 dpf fish show no outersegments in the photoreceptors. A) surface fish photoreceptors and retinal pigmented epithelium (RPE), outer photoreptor segments (OS), inner segments (IS), outer plexiform layer (OPL) and the top part of the inner plexiform layer (INL) can be seen. B) Tinaja cavefish, a pigmented RPE can be seen as well as photoreceptors with missing outer segments. INL with apoptotic cells cells can also be seen. C) Pachon with a degenerating RPE and disorganized OPL. Cell bodies of photoreceptors can be seen and note the presence of melanin in the RPE of surface fish and Tinaja. Scale bar 2um.

## Discussion

*Astyanax* is the only cavefish that can be bred in captivity. The fish are born with eyes, which degenerate during larval development. Here we examined the development of the retina in two parallel lineages of cave *Astyanax*, using immunohistochemistry, TEM and histological staining at 3 days and 7 days post fertilization. We chose these time points because they are before and after the innervation of the optic tectum by RGCs (personal observation). Overall, the neuroretina is similarly laminated with similar cells types in the cave and SF. The main finding is that in the cavefishes the photoreceptor layer is the most disorganized, and that photoreceptor cells are missing outer segments. It is possible for cavefish larvae visual system to function because opsins in the inner segments of photoreceptors could respond to light. If this is the case, we expect remarkable decrease in light sensitivity without the staked discs of the outer segments, and a decrease of resolution in the retinotopic map due to the disorganization of the photoreceptors.

Neurogenesis is generated from the vertebrate Ciliary Marginal Zone (CMZ). The vertebrate eye is structurally conserved, so that eyes of *Astyanax* share the same gross structure as other teleost fish or mammalian eyes; containing a cornea, lens, vitreous, retina, pigmented epithelium, choroid and sclera (Figure 1). It has been well established that the retina grows via a combination of cellular hypertrophy, neurogenesis and intraocular expansion (Ali, 1964; Fernald, 1990; Johns, 1977, 1981; Johns and Easter, 1977; Lyall, 1957; Meyer, 1978; Müller, 1952; Sandy and Blaxter, 1980). Cells that contribute to the increase of the size of the retina are largely born on its rim near the lens, the ciliary marginal zone (CMZ), and creates new tissue concentrically (Amato et al., 2004; Centanin et al., 2011; Cerveny et al., 2012; Harris and Perron, 1998; Hitchcock et al., 2004; Hitchcock and Raymond, 2004; Moshiri et al., 2004; Otteson and Hitchcock, 2003; Raymond et al., 2006; Stenkamp, 2007). In contrast, the CMZ in absent in mammals and has limited neurogenesis of early postnatal birds (Kubota et al., 2002). How precursor cells proliferate, the gene expression, and signaling pathways have been studied extensively in the CMZ (Alunni at al. 2007; Cerveny et al., 2010; El Yakoubi et al., 2012; Kubota et al., 2002, Meyers et al., 2012; Moshiri et al., 2004, Perron and Harris, 2000; Raymond et al., 2006; Reh and Levine, 1998; Stenkamp, 2007; Xue and Harris, 2012). We saw the CMZ in *Astyanax* as a dark border in cresyl violet sections in surface and cavefish and in both time points. This means that *Astyanax* retains neurogenesis capability throughout development including the stages in which the retina is degenerating in cavefish.

Muller cells are radial glia with nuclei in the INL extending processes apically that span the width of the retina, being among the earliest retinal cells to become differentiated (Ramon y Cajal, 1972). These cells provide structural and metabolic support for neurons (Sarthy and Ripps, 2001; Bringmann et al., 2006; Reichenbach and Bringmann, 2013). In both surface and cavefish *Astyanax* we see a basal triangular shape leading to a thin apical processes (Figure 2). We propose that the cell body at these stages are in the basal region, which has not been seen in other fish (Thummel 2008; Goldman 2014; Lenkowski & Raymond 2014). There is evidence that rods are not born in the CMZ but derive from proliferating precursors in the ONL (Ahlbert, 1976; Lyall, 1957; Vilter and Lewin, 1954; Sandy and Blaxter, 1980; Johns and Fernald, 1981). In the zebrafish proliferating Muller cells are the source of rod progenitors (Bernados et al., 2007). In other larval fish studies, such as cichlids and goldfish, the production of cells in the INL form a chain of progenitor cells that move apically into the ONL where they differentiate into rods (Hagedorn and Fernald 1992; Johns, 1982).

In some teleost larvae the retina contains cones but few to no rods, which begin to appear later in life (Blaxter, 1975; Blaxter and Staines, 1970). Here we describe this pattern in *Astyanax*. Cells were positive for ZPR3, that label rhodopsin in the outer segments of the red green double cones as early as 3 dpf. It has been proposed (Evans and Fernald, 1990; Helvik et al., 2001) that the delayed emergence of rods is attributed to changes in the photic environment of the developing fish, as larvae live near the surface and adult fish move down the water column. As ambient light decreases, rods become more important (Müller, 1952). Cones primarily function under different lighting conditions than that of rods, which mediate dim light (scotopic) vision at night while cones mediate bright light (photopic) vision during the day (Dowling, 1987). Both photoreceptor cells have an inner segment, with the nucleus, mitochondrian and an outer segment that contains stacks of discs that contain the molecular machinery for phototransduction. Cone photoreceptors of different subtypes express different opsins that mediate their spectral sensitivity. In turn, the spectral sensitivities of cone photoreceptors determines the fish color vision (Risner et al., 2006). The immuno marker ZPR1 labels arresting3a, which is typically expressed in green and red double cones (Wan and Stenkamp, 2000; Vihtelic and Hyde 2000). In surface *Astyanax* cells are labeled throughout their cytosol, outer and inner segments included, and the peducle of the cone photoreceptor is easily distinguished. In both forms of cave *Astyanax* ZPR1 is seen in what appear to be the inner segments of the photoreceptors. No outer segments were seen and the cells were disorganized in the outermost region of the eye (Figure 9A, 9B). TEM results (Figure 10) confirmed the absence of outer segments and discs at both stages. In another cavefish, namely Sinocyclocheius anophthalmus (Meng et al., 2013) rods and cones have outer segments but smaller in size and disorganized in appearance compared to closely related surface species. It is interesting to note that the lack of outer segments resembles the early stages of retinitis pigmentosa (RP; Milam et al., 1998), where the photoreceptors die after the outer segments shorten and eventually disappear (Milam and Li, 1995; Milan et al., 1996). In contrast to RP, in *Astyanax* the inner segments are disorganized and not a laminated monolayer that eventually degenerates (Milam et al., 1998). RP can be caused by one of 70 mutations in the Rhodopsin targeting pathway (Milam et al., 1996) and it is possible that *Astyanax* share some of these mutations and that in the adult they lose their inner segments. Alternatively, there could be abnormalities in the interactions between photoreceptor and RPE. O’Quin et al. (2013) also suggested comparisons between *Astyanax* and RP and reported cell death via apoptosis in all cell layers. We failed to confirm this with TEM and saw no apoptotic photoreceptors, but we did see apoptotic cells in the ONL, INL, and GCL (Figure 10). Naturally occurring sensory deficits could provide us with insight about Human genetic disorders. In another cavefish, namely *Amblyopsis spelaea*, the histological source of its blindness is yet unknown (aside from sunken, degenerated eyes) and the fish also show deficiencies in hearing, having a shorter frequency range that its surface relatives (Niemiller et al., 2013), reminiscent of human Usher syndrome, where there are deficits on both sensory modalities. The study of the developing and degenerating eye in cavefishes could benefit research of various degenerative human eye diseases such as photoreceptor degeneration by drawing parallels between adaptations and diseases.

## Acknowledgments

We are thankful for Dr. Phillip Barden for the use of the dissecting microscope and Dr. Gal Haspel for comments on the text.

Grant support: NIH R15EY027112

The data that support the findings of this study are available from the corresponding author upon reasonable request.

